# Data Fusion by Matrix Completion for Exposome Target Interaction Prediction

**DOI:** 10.1101/2022.08.24.505125

**Authors:** Kai Wang, Nicole Kim, Maryam Bagherian, Kai Li, Elysia Chou, Justin A. Colacino, Dana C. Dolinoy, Maureen A. Sartor

## Abstract

Human exposure to toxic chemicals presents a huge health burden and disease risk. Key to understanding chemical toxicity is knowledge of the molecular target(s) of the chemicals. Because a comprehensive safety assessment for all chemicals is infeasible due to limited resources, a robust computational method for discovering targets of environmental exposures is a promising direction for public health research. In this study, we implemented a novel matrix completion algorithm named coupled matrix-matrix completion (CMMC) for predicting exposome-target interactions, which exploits the vast amount of accumulated data regarding chemical exposures and their molecular targets. Our approach achieved an AUC of 0.89 on a benchmark dataset generated using data from the Comparative Toxicogenomics Database. Our case study with bisphenol A (BPA) and its analogues shows that CMMC can be used to accurately predict molecular targets of novel chemicals without any prior bioactivity knowledge. Overall, our results demonstrate the feasibility and promise of computational predicting environmental chemical-target interactions to efficiently prioritize chemicals for further study.

## Introduction

Human disease is often not caused by genetic variation^1^ but by environmental factors alone or in combination with genetics; however, the effects of environmental factors on phenotype are not as well studied due to the complexity of environmental exposures^2^. Environmental chemicals enter the human body mainly through three routes of exposure (inhalation, ingestion, or dermal) with the Global Burden of Disease project^3^ estimating that 9 million deaths per year are caused by air, water, and soil pollution alone. However, this number is likely an underestimate due to the fact that many new chemical compounds with uncharacterized toxicological profiles are created and released into our environment annually.

Computational methods are widely used in toxicology research to organize and analyze *in vitro* toxicity and safety assessment data^4,5^. Corresponding methods have also been developed and applied to predict chemical toxicity. Toxicity prediction methods include structure-activity relationship (SAR)^6,7^, read-across^8,9^, and quantitative structure-activity relationship (QSAR)^10–12^. These methods usually focus on using chemical features to predict a binary label for chemicals, such as whether a chemical is a carcinogen or not, or whether the chemical activates certain marker genes or related biological pathways. Thus, the aforementioned methods cannot provide a comprehensive understanding of the biological responses caused by a chemical. Furthermore, these methods do not exploit the vast amount of data accumulated to date for similar chemicals and molecular targets. Therefore, a method that takes advantage of both chemical features and biological knowledge, as well as the known complex relationships among them, could radically improve and expand the scope of chemical toxicity predictions.

The unsolved problem of exposure-target prediction shares similarities with drug-target interactions (DTI) prediction, for which a plethora of computational methods exist, including deep learning, tree-based, network-based, and matrix completion methods. Matrix completion is a task to find the missing values in a partially observed matrix, and has been widely used in DTI prediction. An advantage of matrix completion algorithms compared to other types of methods is they can easily handle big datasets with high sparsity without overfitting^13^. In DTI prediction, a binary value (i.e., 1 or 0) is often used to indicate the interaction between a chemical drug and its molecular targets, because drugs are typically designed with specific molecular targets in mind and thus off-target effects are avoided. In contrast, chemical exposures were not designed to interact with specific gene targets, and therefore they often have broader ranging effects, suggesting a continuous value approach is more appropriate than the typical binary approach used for DTI.

In this study, we take an important step towards the goal of accurate computational exposure toxicity prediction by implementing a new matrix completion algorithm^14^ called coupled matrix-matrix completion (CMMC), which predicts the target genes of environmental chemicals on a broad scale. CMMC falls under the youngest branch of data fusion that is intermediate/partial integration, i.e., it does not necessarily fuse the input data, nor does it develop separate models for each source. Rather, it preserves the structure of the data sources and merges them at the level of a predictive model^15,16^, here CMMC. Our method precisely fits in the next generation blueprint of computational toxicology proposed by the U.S. Environmental Protection Agency to efficiently evaluate the safety of chemicals^17^, by introducing a complementary New Approach Methodology (NAM). CMMC differs from previous matrix completion algorithms based on matrix factorizations and is subject to a Distance Metric Learning (DML) constraint, that is, it directly utilizes supporting information (i.e., coupled matrices about similarities among chemicals and genes) in the main iteration to learn an optimal metric for imputing the missing values (i.e., unknown chemical-gene target pairs). The method also guarantees a unique solution for the optimization problem, a desirable property to have for widespread adoption. We show that CMMC is a powerful untapped tool to predict chemical-gene interactions, which provides vital information for prioritizing further toxicology research.

## Results

### Datasets Used to Predict Gene Targets of Chemical Exposures

The data used as input for this study are from the Comparative Toxicogenomics Database (CTD), a comprehensive, well-maintained publicly available database containing a vast amount of information pertaining to how environmental exposures affects human health, including chemical-gene/protein interactions (Supplementary Table 1). This data included a total of 4,864 chemicals exposures and 201,375 known chemical-gene interaction pairs with 22,606 genes. Three measurements of gene similarities and two measurements of chemical similarities were used. Gene similarities were calculated based on co-occurring GO cellular components (CC), molecular functions (MF), and biological processes (BP) (Supplementary Fig. 1A-C and Supplementary Table 2) while chemical similarity values were calculated using Morgan fingerprints and Topological Torsion fingerprints (Supplementary Fig. 1D, E and Supplementary Table 2). Two different normalization approaches were used to process the interaction values, based on the number of times an interaction was reported (N1) and the number of publications reported for the interaction in the CTD (N2) (see *Methods*). These different normalizations and similarity measurements allowed us to evaluate differences in performance and the sensitivity of the method to differences in set up. Three benchmark datasets ranging in size from 200×400 to 600×800 were generated to evaluate our method and compare its performance with other state-of-the-art coupled matrix completion methods (Fig. 1 and Supplementary Fig. 2).

**Fig. 1.**
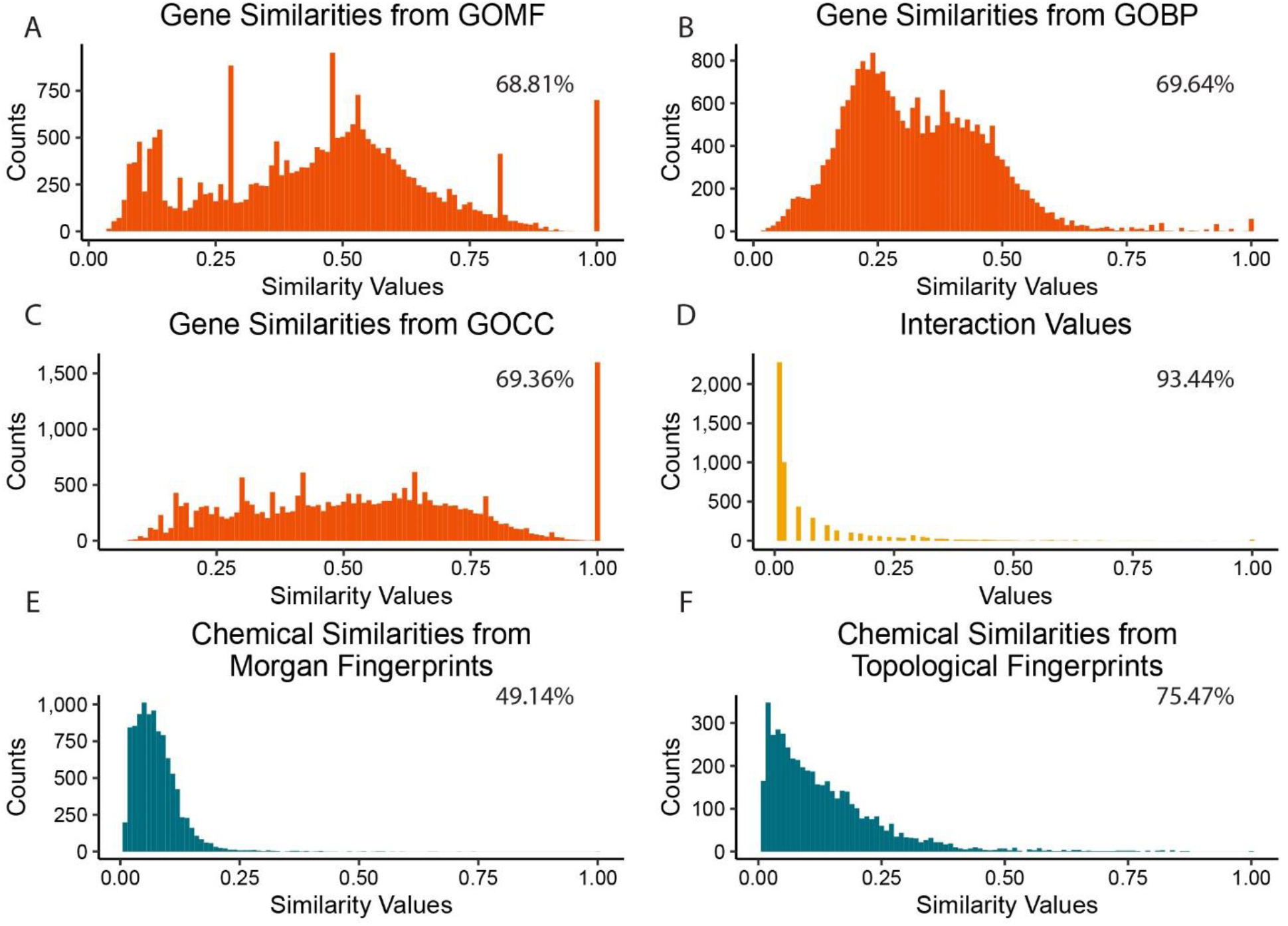
Data distributions for each input matrix for the 200×400 dataset. The sparsity of each matrix is labeled at the right top corner of each histogram. (A-C) gene-gene similarity matrices based on Gene Ontology, (D) the *a priori* known environmental chemical-gene interactions, and (E-F) chemical-chemical similarity matrices based on Morgan and Topological fingerprints, respectively.

### Implementation of CMMC for Chemical Exposure-Gene Target Prediction

In order to implement CMMC for chemical-gene target prediction, two regularization hyperparameters and the number of iterations were optimized, test datasets were generated, and the three sizes of input datasets were created for testing. The hyperparameters and number of iterations were determined using all three different sizes of benchmark datasets (see *Methods*). For performance evaluation, we chose to use the ROC AUC for all three different size of benchmark datasets through selecting true positives and true negatives from our gold standard set. Based on the grid search results, hyperparameters were set to 0.9 for the remainder of the study. We found that different iteration numbers beyond the first had minimal impact on the average AUCs (Supplementary Fig. 3). Thus, optimize run time, two iterations were used in the following analyses.

### Performance Comparison

CMMC was tested and compared with several other state-of-the-art matrix completion methods, namely CMF^18^ (Collaborative Matrix Factorization), GRMF^19^ (Graph-Regularized Matrix Factorization), and WGRMF^19,20^ (Weighted Graph-Regularized Matrix Factorization), all with and without WKNKN^19^ (Weighted K Nearest Knows Neighbors) using the same benchmark datasets for performance testing. For all other methods, parameters were selected as follows: *k* = 100, *λ*_*l*_ = 2, *λ*_*d*_ = 0.125, *λ*_*t*_ = 0.125, and *K* = 5 if WKNKN is used, where k is the number of latent features, *λ*_*l*_, *λ*_*d*_, and *λ*_*t*_ are regularization parameters, and *K* is the number of nearest neighbors. These parameters were optimized through a test run on our benchmark datasets (200×400) using the built-in function of the GRMF package.

The average AUC values calculated based on the N1 interaction values shows CMMC achieved the highest AUCs for all three sizes of input datasets with a maximum average AUC of 0.893 ± 0.012 using the 400×600 dataset (Fig. 2A). However, the AUC values for the different sized matrices were not significantly different. The average AUCs across different combinations of coupled matrices with the ten different testing datasets for 400×600 input matrix improved from CMF (0.824 ± 0.014) to GRMF (0.836 ± 0.017) to WGRMF (0.843 ± 0.014) with a binary conversion cutoff equals of 0.05. WKNKN further improved the AUC values for these three methods (Fig. 2A and 2B). Results with other binary conversion cutoffs confirmed that CMMC maintained its superior performance in all cases evaluated (Supplementary Table 3), and confirmed that a small binary conversion cutoff is optimal for CMF, GRMF, and WGRMF.

**Fig. 2.**
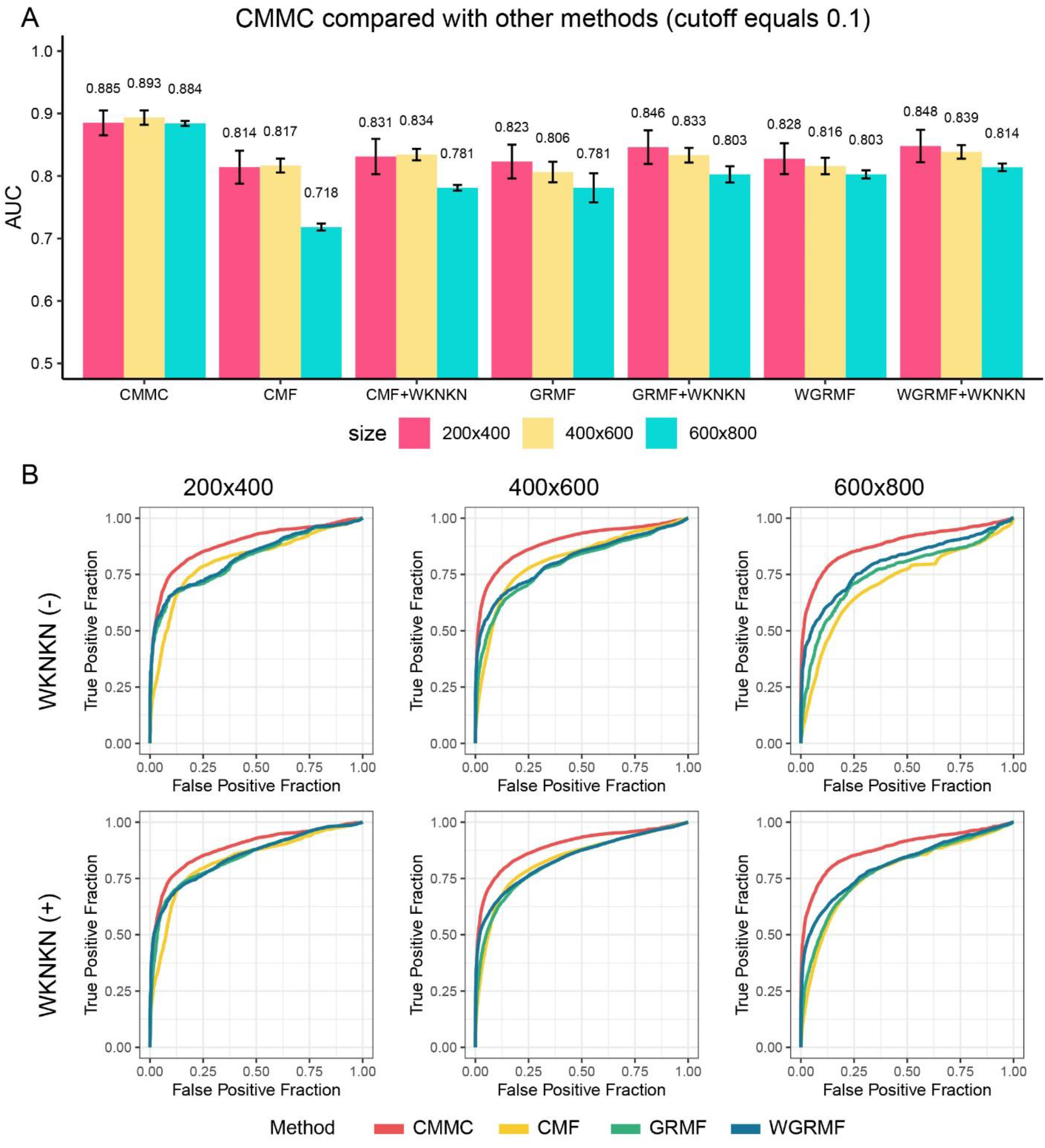
Performance comparison of CMMC with alternative methods. (A) Average AUCs of the methods using the N1 interaction values (see Methods). Binary conversion cutoff is 0.1 for CMF, GRMF, and WGRMF. (B) ROC curves of different datasets (with and without WKNKN for the alternative methods).

Surprisingly, all other methods besides CMMC showed a consistent decrease in performance moving from smaller to larger input matrices, whereas CMMC demonstrated the most consistent performance. It is possible this result could be explained by the higher sparsity level in the larger matrices, a characteristic to which CMMC has been shown to be robust. Comparing the results between N1 and N2 for CMMC, N1 normalization resulted in better performance across all three different sizes of input data, with average AUC values near 0.89 compared to 0.84 for N2. Based on these results, the following case studies were performed using CMMC with N1.

### Case study

We selected the very well-studied endocrine disrupting chemical bisphenol A (BPA) and its closely related but lesser studied analogs BPB, BPF, and BPS, for our case study. These were chosen due to the fact that BPB, BPF, and BPS are often used as alternatives for BPA in manufacturing now that BPA has been banned in many products and is banned completely in some countries^21^. However, given the similarity in BPA’s chemical structure compared to the analogs and based on preliminary toxicology and epidemiology studies in the analogs, negative health effects could also be caused by exposure to the alternatives. For each of the four chemicals, we generated 200 different datasets, each containing a randomly selected additional 299 chemicals and 500 genes from the total 5000 chemicals and 10,000 genes used for this study, respectively. This experimental design allowed us to obtain ∼10 prediction for each chemical-gene pair with fast overall run time (∼30 minutes). All previously known interactions of the case study chemical in the 200 different datasets were removed from the main matrix (M_EG_) to mimic a realistic scenario where one does not have any prior bioactivity knowledge of a new chemical. After imputing all 200 datasets for a chemical, the average imputed interaction values across all six runs (two chemical-chemical similarities by three gene-gene similarities) and 200 datasets were calculated for further analysis. Bioactivity data for BPA, BPB, BPF, and BPS from CompTox Chemical Dashboard and the corresponding GO terms from CTD were used to validate the predicted genes for each chemical (Supplementary Table 4 and 5). Genes not included in this study were removed from further analysis. A large, imputed interaction value in our results indicates a strong, high-confident interaction. Thus, for each chemical, genes were ordered by the imputed interaction values in descending order. Genes with active bioactivity responses (“active genes”) of the four chemicals from the CompTox Chemical Dashboard were compared to our results (Fig. 3). For BPA, 20 of the 54 active genes from CompTox were predicted in the top 100 interactive genes in our results, a 37 fold enrichment over what would be expected by chance. While for BPB, BPF, and BPS, there were 23/74, 10/18, and 7/12 active genes predicted in the top 100 interactive genes, respectively (fold enrichments of 31, 55, and 58; Fig. 3A). Among these genes, four genes were identified as common target genes across these four chemicals, i.e., *ESR1, ESR2, AR*, and *NR1I2*. Furthermore, around 50% of the active genes of BPA and BPB and 80% active genes of BPF and BPS were predicted as top 1,000 genes, respectively (Supplementary Table 4). In the CompTox Chemical Dashboard, seven genes were identified as active genes for all four of these chemicals (*ESR1, ESR2, AR, NR1I2, PGR, PPARG*, and *PPARD*). Among these genes, all six of them used in our study, i.e., *ESR1, ESR2, AR, NR1I2, PGR, PPARG*, were predicted to be in the top 1,000 interacting genes for all four chemicals in our results (Fig. 3B). Since the 10,000 gene list was randomly selected, *PPARD* was not included in the total gene list.

**Fig. 3.**
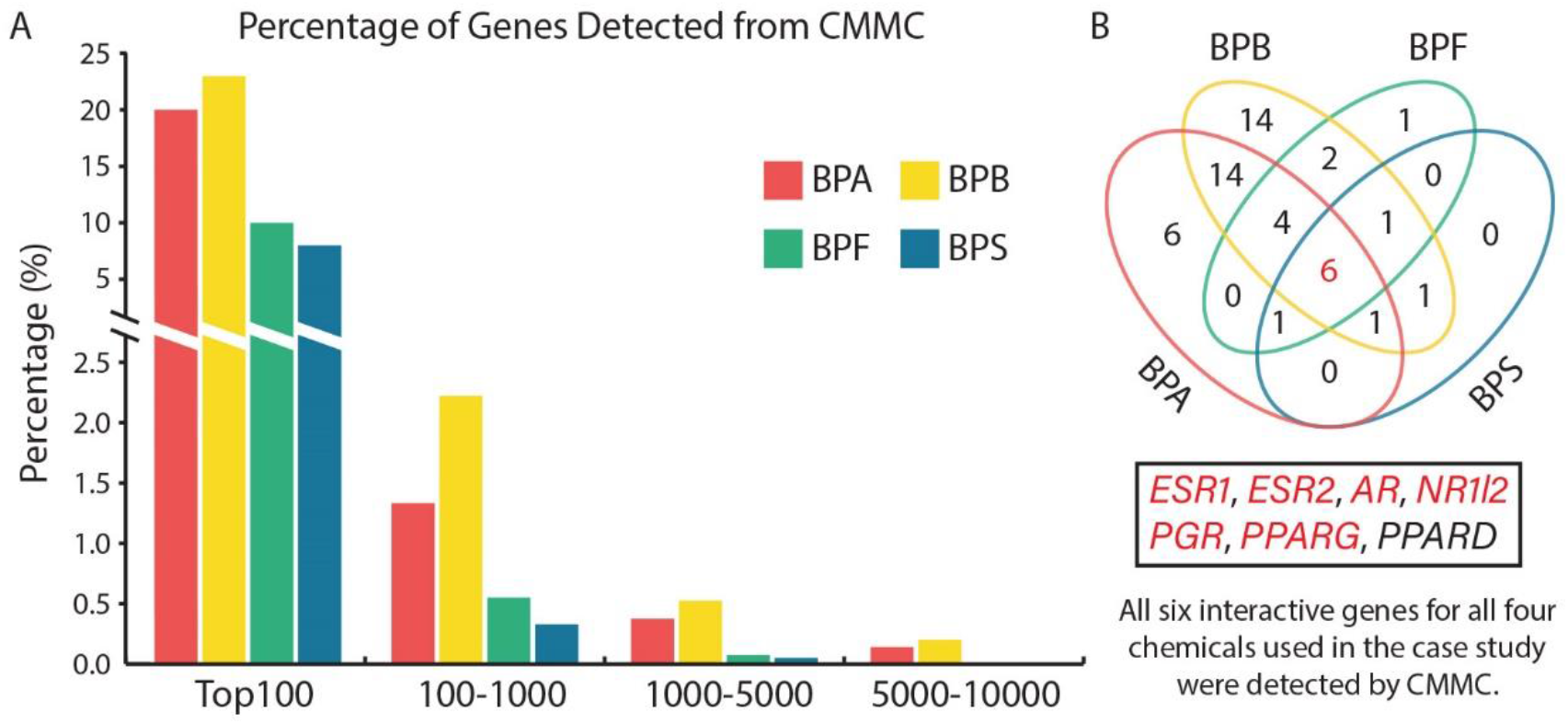
Results validation with CompTox bioactivity data. (A) Percentage of CompTox interacting genes detected by CMMC. (B) Venn diagram of the top 1000 genes corresponding to (A). All six of the seven interactive genes for all four chemicals from CompTox that were used in the case study were also detected as interactive genes in our results.

To further investigate our results, the top 1,000 interacting genes of each tested chemical were used to identify the related GO terms through gene set enrichment testing. After removing GO terms containing more than 500 genes and filtering out GO terms with an FDR larger than 0.05, we found 401, 394, 343, and 396 significant GO terms for BPA, BPB, BPF, and BPS, respectively (Supplementary Table 5). Strikingly, *Response to Estradiol* (GO:0032355), *Positive regulation of ERK1 and ERK2 cascade* (GO:0070374), and *Cellular response to tumor necrosis factor* (GO:0071356) were ranked in the top 5 significant GO terms for all four chemicals. Since BPA is an endocrine-disrupting chemical^22^, hormone related GO terms were extracted from the significant GO term lists (Fig. 4A). Most hormone-related GO terms were significantly enriched for all four chemicals, including *Response to Estradiol* (GO: 0032355), *Response to Estrogen* (GO: 0043627), *Estrogen Metabolic Process* (GO: 0008210), and *Estrogen Receptor Binding* (GO:0030331). However, *Insulin receptor signaling pathway* (GO:0008286) and *Estrogen 16-Alpha-Hydroxylase Activity* (GO:0101020) were only significantly enriched for BPB while *Insulin-like growth factor I binding* (GO:0031994) and *Positive Regulation Of Insulin-Like Growth Factor Receptor Signaling Pathway* (GO:0043568) were only significantly enriched for BPA.

**Fig. 4.**
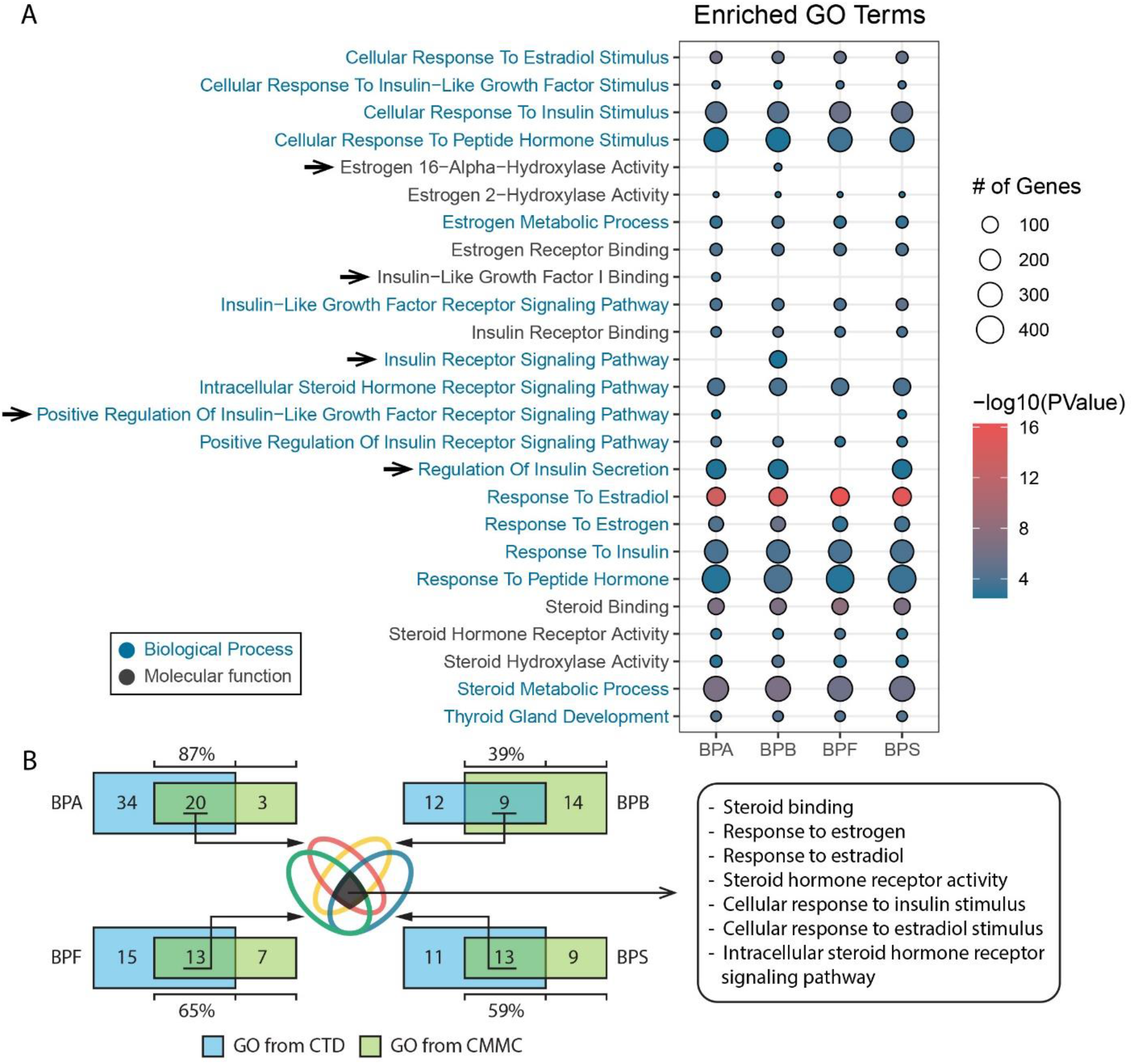
(A) Hormone-related significantly enriched GO terms from gene set enrichment test (Top 1000 interactive genes from the CMMC results used as input gene list). Arrows point to the GO terms that differ by chemical. (B) Overlap between GO terms detected by genes from CMMC prediction and genes directly from the CTD.

For comparison, genes known to interact with these four chemicals were downloaded from CTD and gene set enrichment testing was used to find the significant hormone-related GO terms. Because there are more than 5,000 interacting genes in CTD for BPA and BPS, only the top 1,000 interacting genes were used. For BPB and BPF, all genes in CTD (115 genes for BPB and 144 genes for BPF) were used for gene set enrichment testing. For BPA, 54 hormone related GO terms were significantly enriched from the CTD genes while 23 hormone related GO terms were significantly enriched from the top 1000 genes predicted by CMMC (Supplementary Fig. 4). 87% (20/23) of these hormone related GO terms from the top 1000 genes predicted by CMMC were also enriched in GO terms from CTD genes. This number is 39% (9/23), 65% (13/20), and 59% (13/22) for BPB, BPF, and BPS, respectively (Fig. 4B and Supplementary Fig. 5-7). Among the significantly enriched GO terms using genes from CTD and CMMC, six of them were enriched for all four chemicals (Fig. 4B). CMMC is able to find interacting genes that distinguish the four chemicals, in spite of their similar structures. For instance, *GPER1* was only predicted as a top target gene of BPB^23^. Among the CompTox active genes of these four chemicals, 2 of the 9 BPA unique active genes and 8 of the 26 BPB unique active genes were predicted by CMMC as top target genes (Supplementary Fig. 8).

## Discussion

The U.S. Environmental Protection Agency (EPA) is strongly encouraging the development of New Approach Methodologies (NAMs), which include *in silico* methods to assess potential health impacts of chemical exposures^5,24^, for future toxicology research. In this paper, we illustrated the feasibility of predicting gene targets of novel or understudied chemical human exposures on a large-scale. The significance of this to the field of public health is great given the high global burden of disease and mortality from environmental exposures, the large number of unstudied novel chemicals being produced each year, and the inability to adequately evaluate each chemical with current funding levels. Our new method fills a current major hurdle in computationally prioritizing chemicals for further analysis of their toxic potential. Once the target genes and their predictive scores are calculated, methods already in use such as ProTox-II^25^ and others can be employed to predict the toxicity, target tissues, phenotype, and potential disease risk by incorporation of additional data.

Data fusion by representing data from multiple sources as a collection of matrices is an effective way to jointly analyze data together with supporting information in the context of additional constraints. Matrix completion, as an approach in data fusion, is a prediction method designed to find missing values from a partially observed matrix, where the input is usually a highly sparse matrix. In general, missing values are imputed by using the low-rank matrix of the known values. However, it is extremely difficult to predict the missing values of a new row or column item when there is no relevant prior knowledge. Therefore, matrix completion methods that incorporate supporting information in the form of coupled matrices represent a significant step forward, and have gained traction by showing improved performance in recent years^26,27^. In this study, we created and tested a new implementation of the recently developed CMMC for predicting environmental chemical-gene interactions. The environmental chemical-gene pairs used in this study from the CTD.

We compared the performance of our CMMC implementation for predicting environmental chemical-gene interactions, with three related, published methods (i.e., CMF, GRMF, and WGRMF). Additionally, WKNKN, a preprocessing method for estimating the interaction likelihoods of unknown interactions based on known neighbors, was utilized for the three aforementioned methods in the performance comparison. We chose these three methods because they also permit the use of two coupled supporting matrices as input for prediction. These three methods were originally developed for predicting DTI with a binary interaction matrix, so we used three different cutoffs (i.e., 0.05, 0.1, and 0.2) to convert the interaction values into binary values for input for other methods. The performance of these three methods gradually improved from CMF to WGRMF. Furthermore, WKNKN additionally improved the performance of these three methods. This result is consistent with the previous study that CMF obtained the worst performance^19^. Considering the fact that our nonbinary interaction values are highly skewed towards the left, it is not surprising to see improved performance with smaller binary conversion cutoffs.

BPA and three of its analogues were selected to perform a case study, namely BPB, BPF, and BPS. We chose BPA because it is a well-studied chemical with abundant bioactivity records for validating our results. BPB, BPF, and BPS are chemicals with similar structures and are used as substitutes of BPA in industrial production. These four chemicals have two phenol groups connected through a linking group. Considering the chemical similarities among these four chemicals (Supplementary Fig. 9), we expected similar but not identical prediction results. More than half of the interacting genes of these four chemicals from the CompTox database are identified in our results as top interacting genes. Since CompTox is a data source independent of CTD, our results are independently validated. All known interaction values were removed in the case study for each chemical in the imputation, so this result suggests that our method can be used to effectively identify target genes for novel chemicals. BPA belongs to the endocrine disrupting class of chemicals, and low dose BPA can impact reproduction, the immune system, and is related to hormone-dependent cancers^22^. Previous studies have shown that some nuclear receptors are target genes of BPA, such as androgen receptor^28^ (*AR*), estrogen receptor 1 (*ESR1*), estrogen receptor 2^29^ (*ESR2*), and aryl hydrocarbon receptor^30^ (*AHR*). In our results, all of the aforementioned four genes were predicted as top 100 interactive genes for BPA. Except for *AHR*, the other three genes were also predicted by CMMC as top interactive genes for BPB, BPF, and BPS (Supplementary Table 4). This indicates that the three analogue may have some of the same adverse biological effects as BPA^31^. Furthermore, estrogen and other hormone-related GO terms identified from our gene lists of these four chemicals are very similar to each other and are consistent with the fact that these chemicals are endocrine disruptors and are able to mimic estrogen^32^. The relationship between BPA and type 2 diabetes has been well studied in the last decade^33,34^. A recent longitudinal study provide new evidence that not only the BPA but also BPS are associated with type 2 diabetes^35,36^. In our results, several insulin related GO terms are significantly enriched from our gene list of all four chemicals. Thus, the CMMC results prioritize the biological effects of BPF and BPB on type 2 diabetes for future examination.

The supporting information in the two coupled similarity matrices is critical for any prediction method to find chemical-gene interactions for a novel chemical without any prior knowledge. The input information is very flexible, thus it will be interesting to test alternative similarity metrics, such as gene-gene sequence or expression similarities instead of or in combination with gene-gene similarity based on GO term membership. Using this coupled matrix information with an optimized rather than fixed similarity measure represents a significant strength of CMMC. Furthermore, CMMC is not a black box method. By comparing the similarities of the chemicals being predicted with other known chemicals, the similarities of the known predicted genes for the most similar chemicals with the genes predicted for the new chemicals, and the similarities among the top-ranked genes, we can gain direct insight into the causes for each prediction. For example, G protein-coupled estrogen receptor 1 (*GPER1*) was only predicted in the top 100 target genes of BPB by CMMC in our case study. This is consistent with a previous study that BPB and BPAF (Bisphenol AF) have an approximate 9-fold higher binding affinity to *GPER1* than BPA^23^. BPAF is an analogue of BPA in which the two methyl groups are replaced with trifluoromethyl groups. Although all five of these chemicals are very similar, BPB and BPAF are more similar to each other in terms of adding/replacing chemical groups to the linking group of BPA. This can partially explain why only BPB among the four chemicals was predicted to target *GPER1*, that is, it learned from the similarity with BPAF. This information is very useful for post-hoc analyses and follow-up studies and would not be so easily determined with a deep learning approach.

This study has multiple limitations. Currently, we did not differentiate the chemical-gene interaction categories that are provided in the CTD. Although most interactions are labeled as changes in RNA expression changes (52.6%), there are 132 different interaction categories listed. Appropriately incorporating chemical-gene interaction categories into the prediction could be advantageous for future studies. Identifying the target genes of a novel chemical is key but is just one step in understanding the potential health effects from an environmental exposure. Knowing the target tissues, considering the dosage level, and linking the molecular response to diseases are all downstream steps that are not considered in this paper. However, CMMC combined with our data normalization strategy has great potential to form a prioritization pipeline for discovering potential molecular targets of novel chemicals without any priori bioactivity knowledge. Future studies could potentially incorporate tissue information, dose information, and/or interaction category information as an additional dimension in the interaction matrix to form a partially observed tensor, thus expanding the CMMC to CMTC (coupled matrix-tensor completion) for missing value imputation.

We envision that a matrix completion approach like CMMC that incorporates information from already known interactions of similar chemicals and similarities among their gene targets will provide complementary information to the currently used toxicity prediction methods, such as QSAR. Used in conjunction, they have the potential to significantly improve efforts to prioritize chemical exposures for further study.

## Methods

### Overview of the CMMC Algorithm

Matrix completion is an ideal approach for our goal due to the great deal of information already accumulated and available in databases about the genes affected by chemical exposures. CMMC, in particular, is attractive because it not only takes advantage of the known chemical-gene interactions, but also the hidden feature relationships among chemicals and among genes. CMMC was originally developed for predicting DTI with a binary input. As opposed to other coupled matrix completion methods, which use a fixed distance metric, such as *Euclidean*, CMMC uses DML in order to learn the metric from the data itself^14^.

A low-rank matrix completion for an incomplete unobserved/under-observed matrix X can be formulated as:

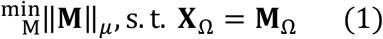

where Ω denotes an index set and *μ* is a proper metric to be determined. In other words, given an incomplete unobserved/under-observed matrix **X**, the goal is to find matrix **M** whose rank is minimum, and it has all the known entries in **X**. Most methods for matrix (or tensor) completion rely upon the choice of a fixed metric, for instance the Euclidean or nuclear norm. If there is a high correlation between the rows/columns in a matrix, then a different metric given by the data itself could be adopted. This is part of a well-known branch in machine learning known as Distance Metric Learning that aims to learn distances from the data^37^. When working with problems including data points in n-dimensional space, ℝ^*n*^, *Mahalanobis distance*, as the inverse of covariance matrix of the data, learns a metric induced by the data points itself. As proposed in Bagherian M, et al^14^, CMMC method development is heavily based on the algebraic property of symmetric matrices which can be thought of elements of *General Linear* (GL) and *Special Linear* (SL) *Groups*. These linear groups belong to a class, called reductive groups^38^. Norm minimization problems when put in the form of reductive groups (with a proper group action) can in fact obtain a unique minimum (up to an orbit) using the algebraic results given in *Kempf-Ness theorem*^38^. *Mahalanobis distance* is equivalent to *Euclidean distance* after a linear change of coordinate and normalization with respect to covariance matrix. The *Kempf-Ness theorem* shows that there is essentially a *unique* change of coordinates that is optimal in a certain sense. CMMC falls under the youngest branch of data fusion that is intermediate/partial integration, i.e., it does not necessarily fuse the input data, nor does it develop separate models for each source. Rather, it preserves the structure of the data sources and merges them at the level of a predictive model^15,16^.

Let data matrix **X** ∈ ℝ^*n×m*^ be given, where to any of its modes, *n* and *m*, auxiliary information in form of symmetric matrices is present, i.e., adjacent to the mode *n*, there exists an *n* × *n* square symmetric matrix, called **M**_*nn*_ and the same for the mode *m* and matrix **M**_*mm*_. These two coupled/auxiliary matrices can be thought of regularization terms of the matrix completion problem.

The goal is to find completed observed matrix, **Y** ∈ ℝ^n×m^ such that the known entries of **X** and **Y** match. To this end, through the following optimization problem, we first learn two matrices **S**_*nn*_ and **S**_*mm*_ using the given side information, **M**_*nn*_ and **M**_*mm*_, and use them to learn the low rank completed version of **X**, i.e., **Y**. Therefore, the optimization problem to be minimized can be outlined as:

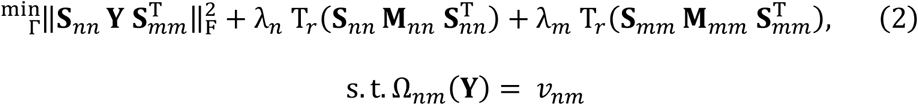

over the optimization set Γ = {**S**nn ∈ **S**L_*n*_, **S**_*mm*_ ∈ **S**L_*m*_, **Y** ∈ ℝ^*n×m*^} and for some regularization parameters *λ*_*n*_, *λ*_*m*_. The optimization problem given in Eq. 2 is subject to a vector constraint *v*_*nm*_. If the known entries of **X** (or **Y** since the known entries match) are at positions

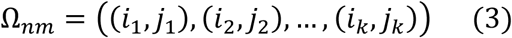

this constraint can be written as

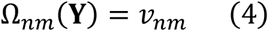

where *v*_*nm*_ ∈ ℝ^*k*^ is some fixed vector consisting of only known then fixed entries. If it happens that the auxiliary matrices **M**_*nn*_ and **M**_*mm*_ are under-observed, the vector constraint may be imposed over these two matrices as a *pre-processing* step. The additional constraints may be written as:

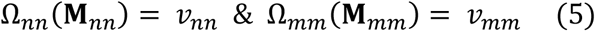

It is noteworthy that without loss of generality one may perform an extra normalization step with respect to the non-zero determinant of a given symmetric matrix in order for it to belong to the special linear group, SL. Additional constraint, such as positive-definite constraint, or other convex regularization term may also be integrated to the main objective function given in Eq. 2, should they be required.

Given the objective function in Eq. (2), M_EG_ corresponds to the matrix, **Y**, which itself is formed by known values only of initial matrix of gene-chemical interaction. Square symmetric matrices M_EE_ and M_GG_ correspond to **M**_*nn*_ and **M**_*mm*_, respectively (Fig. 5). The goal is to learn matrices **S**_*nn*_ and **S**_*mm*_, using DML, through which the missing values of M_EG_ are predicted. M_EE_ and M_GG_ can be thought of as regularization matrices, and two regularization hyperparameters, *λ*_*X*_ and *λ*_*Y*_ as *λ*_*n*_ and *λ*_*m*_ in Eq (2), are applied to scale and add weights to the diagonal for preprocessing M_EE_ and M_GG_, respectively. Observed items (i.e., known chemical-gene interactions) in M_EG_ are used as constraints when minimizing the objective function. Two output matrices (B_1_ and B_2_) are initialized as identity matrices that are the same size as the two coupled input matrices. The comprehensive mathematical description of CMMC can be found in Bagherian M, et al^14^.

**Fig. 5.**
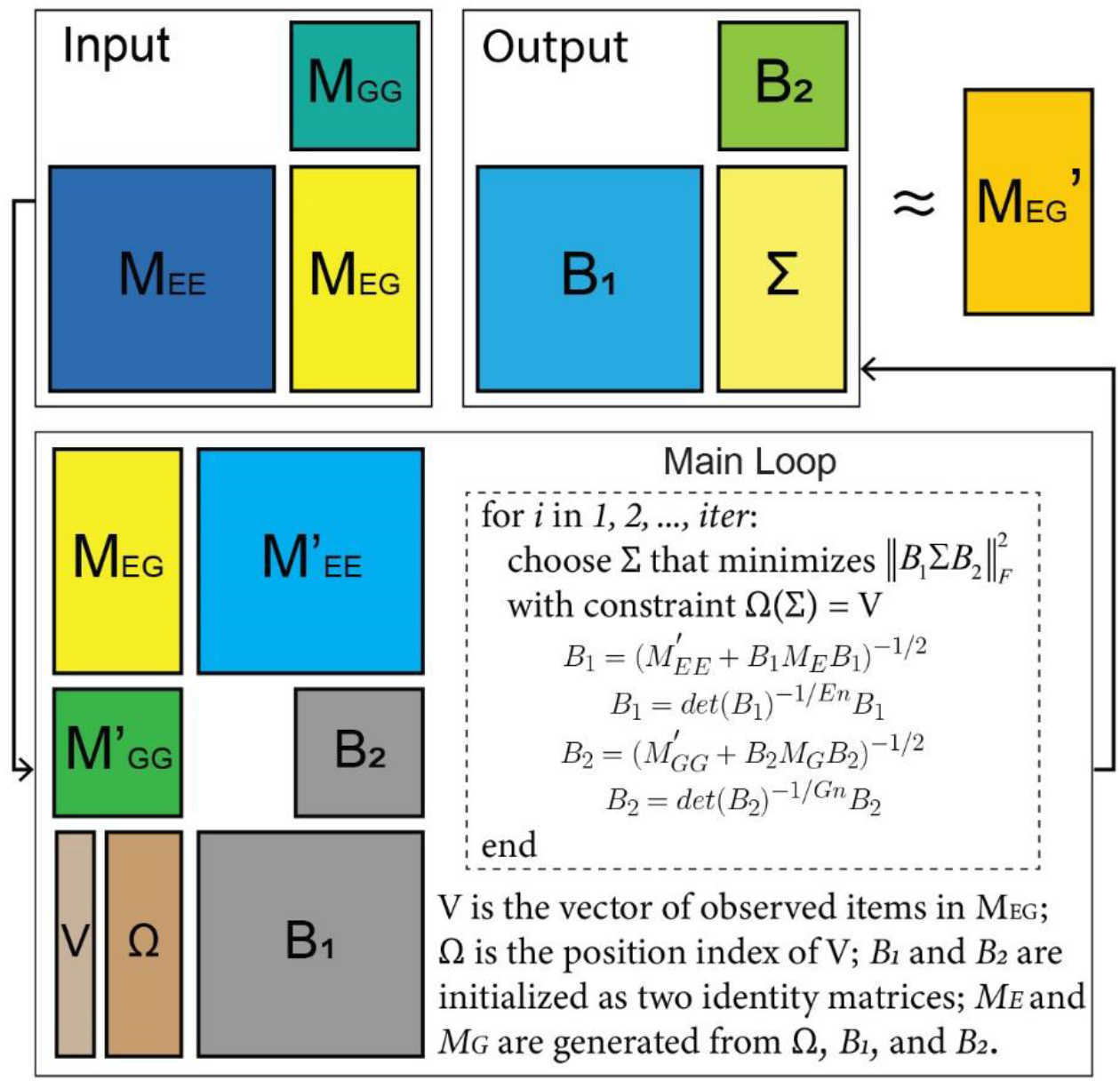
Illustration of the coupled matrix-matrix completion algorithm. Three input matrices are M_EE_, M_GG_, and M_EG_. M’_EE_ and M’_GG_ are the preprocessed coupled matrices with the two hyperparameters.

### Chemical-gene Interaction Dataset and Normalization

We downloaded chemical-gene interaction data from the Comparative Toxicogenomics Database (CTD v16202)^39^. Human-related interactions were extracted and preprocessed for this study. In total, the CTD reported 871,112 chemical-gene interactions documented in humans. Since this study is focused on environmental exposure chemicals, any drugs listed in the CTD were excluded (Supplementary Fig. 10). After the drug filtering, 4,864 chemicals and 22,606 genes with 201,375 known interactions from human studies were included for further data normalization, resulting in a sparsity level of 99.8% for the total chemical-gene matrix. For this study, we calculated an *a priori* interaction value for each known chemical-gene pair and normalized these values in two ways without considering the different interaction categories. The first normalization method (N1) uses the number of occurrences of a chemical-gene pair in the dataset. The second method (N2) uses the number of unique PubMed IDs^40^ listed for a chemical-gene pair (same chemical-gene pairs could be reported in different publications). Since approximately 78% of the chemical-gene pairs had only one interaction record in the dataset, the resulting distribution was highly skewed towards zero. A log transformation was therefore applied to both N1 and N2 interaction values to reduce the skewness of their respective distributions. Thus, for a specific chemical-gene pair, there were two values to represent the interactions. Finally, interaction values were scaled to be in the range of 0 to 1 (Supplementary Table 1).

### Chemical-chemical Similarities

Two widely used chemical fingerprints^41^ were applied to calculate the chemical-chemical similarities, i.e., Morgan fingerprints and Topological Torsion fingerprints. The SMILES^42^ (Simplified Molecular-Input Line-Entry System) string was used for the aforementioned two fingerprint calculations. The CIRpy (v1.0.2) and pubchempy (v1.0.4) Python packages were used to extract the SMILES strings from the PubChem database for all chemicals in our dataset. Chemical-chemical similarities were calculated using the RDKit Python package^43^ (v2020.09.1). Chemical similarities calculated using Morgan fingerprints are highly skewed towards zero with a clear peak around 0.1, while chemical similarities calculated using Topological Torsion fingerprints were also skewed towards zero, but with a data peak around 0.02 with a small kurtosis (Supplementary Fig. 1D, E).

### Gene-gene Similarities

There are many ways to calculate gene-gene similarities. Pair-wise sequence alignment algorithms are widely used for calculating gene similarities based on either DNA/RNA sequences or protein sequences of genes. However, these methods cannot precisely reflect the biological similarities among genes. Thus, instead of using a sequence alignment-based method for calculating the gene-gene similarities, we used a Gene Ontology^44^ (GO) based method called GOSemSim^45^ to measure the gene-gene similarities. Since there are three domains of GO annotation, i.e., cellular component (CC), molecular function (MF), and biological process (BP), three gene similarity matrices were generated for this study. Many gene pairs were assigned a similarity value of 1 based on GOCC and GMMF, while GOBP gene similarities were skewed towards to zero with a peak around 0.2 (Supplementary Fig. 1A-C).

### Generation of Benchmark Datasets

To comprehensively evaluate CMMC on environmental chemical-gene interaction datasets, we generated three different sizes of input matrices with corresponding coupled matrices: 200 chemicals by 400 genes (200×400), 400 chemicals by 600 genes (400×600), and 600 chemicals by 800 genes (600×800). To make our nonbinary interaction datasets comparable to previously published methods used for binary interaction predictions, we manually created two gold standard datasets to use as true positive and true negative sets for performance testing. As described above, for each chemical-gene pair, two different interaction values, N1 and N2, were calculated. For the true positive dataset, only the most confident chemical-gene pairs, i.e., those that have more than 20 interaction records (not unique PubMed IDs) were selected, which resulted in 462 chemical-gene pairs in the true positive dataset. To generate the true negative dataset, chemicals were sorted by the number genes they interact with to select for the most well-studied chemicals. Then, the top 20 chemicals with the greatest number of interacting genes were selected as chemicals for the negative dataset. In this manner, we generated a chemical-gene pair list with the 20 chemicals and the genes having no interaction evidence with these 20 chemicals as a true negative dataset.

To perform robust performance testing, for all three different sizes of input matrices, 10 different sets of randomly selected chemical-gene pairs from the positive and negative datasets were used to generate 10 different input interaction matrices separately for a 10-fold cross validation. The number of chemical-gene pairs for each fold took up 0.25% of the total size of the input interaction matrix. For instance, 0.25% of a 200×400 interaction matrix equals 200. Thus, 100 chemical-gene pairs were randomly selected from both the positive and negative data sets separately to form a total 200 chemical-gene pairs for one-fold. Since there are only 462 chemical-gene pairs in the positive dataset, all of these interactions were used in each of the ten folds of the 600×800 datasets. However, the 600 chemical-gene pairs in the true negative set were different in each fold for testing. Both types of normalized interaction values (i.e., N1 and N2) were used to generate the benchmark datasets, separately.

### Parameter Optimizations

A small test on the 200×400 dataset found that the performance of CMMC was not greatly impacted by different iteration numbers (i.e., 1, 2, and 3) with hyperparameters larger than 1 (data not shown). Thus, a grid search with five values (i.e., 0.05, 0.1, 0.3, 0.5, and 0.9) was used to find an optimal hyperparameter pair for the analysis in this study. To reduce the run time of CMMC on hyperparameter selection, the iteration number was set to 1 for all calculations. Because we generated two different types of chemical similarities, three types of gene similarities, and two different types of chemical-gene interaction values for this study, there were a total of twelve different scenarios for which we tuned the hyperparameters. In addition, for each of the three sizes of input matrix and corresponding coupled matrices, ten different testing datasets were generated for a 10-fold cross validation. Thus, for each size of input dataset with one pair of hyperparameters, AUCs were calculated and averaged from 120 runs (2×3×2×10). The average AUC values increased with large parameters in all three sizes of input data (Supplementary Fig. 11).

After selecting the optimal hyperparameters, we sought to obtain an optimal iteration number for CMMC on exposome target predictions. When applying this algorithm to a binary DTI matrix, this algorithm converged in less than 5 iterations. The benchmark dataset was used to search for an optimal iteration number from 1 to 6. The average AUC values were calculated using the same procedure described above.

### Implementation of Other State-of-the-Art Methods

Similar matrix completion methods are broadly used in the related problem of DTI predictions. Thus, we compared our method to other widely used matrix completion methods to test the performance with respect to the area under the ROC curve (AUC). These methods are CMF^18^, GRMF^19^, and WGRMF^19,20^. These methods were selected for performance comparison because they also require supporting information to impute missing values in the interaction matrix. WKNKN^19^ is a preprocessing method that can be used to estimate the interaction likelihoods for missing values in an interaction matrix. With and without WKNKN, our method was compared to other six methods, i.e., CMF, CMF+WKNKN, GRMF, GRMF+WKNKN, WGRMF, and WGRMF+WKNKN. The same testing datasets were used for performance comparison testing. Since the aforementioned three methods were developed for imputing missing values in a binary interaction matrix, our nonbinary interaction matrices were converted to binary matrices for these methods by applying three cutoff values separately, i.e., 0.05, 0.1, and 0.2. Any interaction values larger than the cutoff were replaced by 1 while those smaller than the cutoff were replaced by 0. Then, the area under the ROC curve (AUC) values were calculated for performance comparison.

### Runtime of CMMC

Since our implementation of CMMC is in C++ and the other methods are implemented in MATLAB, we felt it would be unfair to compare the run times. However, the elapsed real time of CMMC (excluding any data preparation steps) on the benchmark datasets were very reasonable (Supplementary Fig. 12). The average run time for a 600×800 input matrix it is 10.1 ± 5.4 seconds. The run times of the alternative methods implemented in MATLAB were on the order of 10 to 25 minutes for processing the benchmark datasets.

### Generation of Case-study Datasets

The ultimate goal of this study is to predict potential gene targets of novel chemicals without any prior knowledge of biological effects. Thus, to test our method on a real-life scenario, we generated a series of datasets for BPA, BPB, BPF, and BPS. To perform a comprehensive case study, we generated 200 different datasets, each with 299 randomly selected chemicals (300 chemicals in total when including the case chemical) and 500 genes from a total 5000 chemicals and 10,000 genes for each case-study chemical, respectively. Since there are 4,864 chemicals after filtering drugs from the CTD dataset, chemicals from US EPA’s Toxic Substances Control Act Chemical Substance Inventory (TSCA Inventory) were used to form a chemical list with 5000 chemicals for this case study. To mimic a real-life scenario with missing information, rather than only use a small handful of genes, we used a substantial portion of all genes (10,000) randomly selected from the total 22,606 genes without specifically including all interactive genes of these four chemicals. The complete gene list and chemical list can be found in Supplementary Table 6. In this way, interactions between each case-study chemical and 10,000 genes were imputed 10 times, and the resulting prediction averaged. In all 200 chemical-gene interaction matrices for each case-study chemical, all known interaction values were removed to mimic a real-life scenario. Bioactivity data of BPA, BPB, BPF, and BPS were downloaded from the CompTox Chemical Dashboard for validating our prediction results. The DAVID^46^ (Database for Annotation, Visualization and Integrated Discovery) web service was used to identify related GO terms using the genes predicted from CMMC and genes from CTD of the four chemicals in case study, respectively. GO term sets containing more than 500 genes were filtered out for further analysis. An FDR less than 0.05 was used for selecting significant GO terms. The R Bioconductor package org.Hs.eg.db^47^ (v3.14.0) was used for corresponding GO annotations.

## Data availability

The authors declare that the data supporting the findings of this study are available within the paper and its supplementary information files. The source code of CMMC and the benchmark datasets are publicly available at https://github.com/sartorlab/CMMC-on-Exposome-Prediction.

## Acknowledgments

We thank Dr. Kayvan Najarian and Dr. Renaid Kim for their professional help of this study, and we also thank the three anonymous reviewers for reviewing this manuscript. Support for this research was provided by NIH/NIEHS grants P30ES017885, R35ES031686, and R01ES028802.

## Author contributions

M.S. and K.W. conceived of the study. K.W. implemented the algorithm. K.W., N.K., K.L., and E.S. collected the data and performed the data analysis. K.W. and M.B. wrote the manuscript and all authors edited and approved the manuscript.

