## Supplementary material for "Data Fusion by Matrix Completion for Exposome Target Interaction Prediction": Supplementary.Information.docx

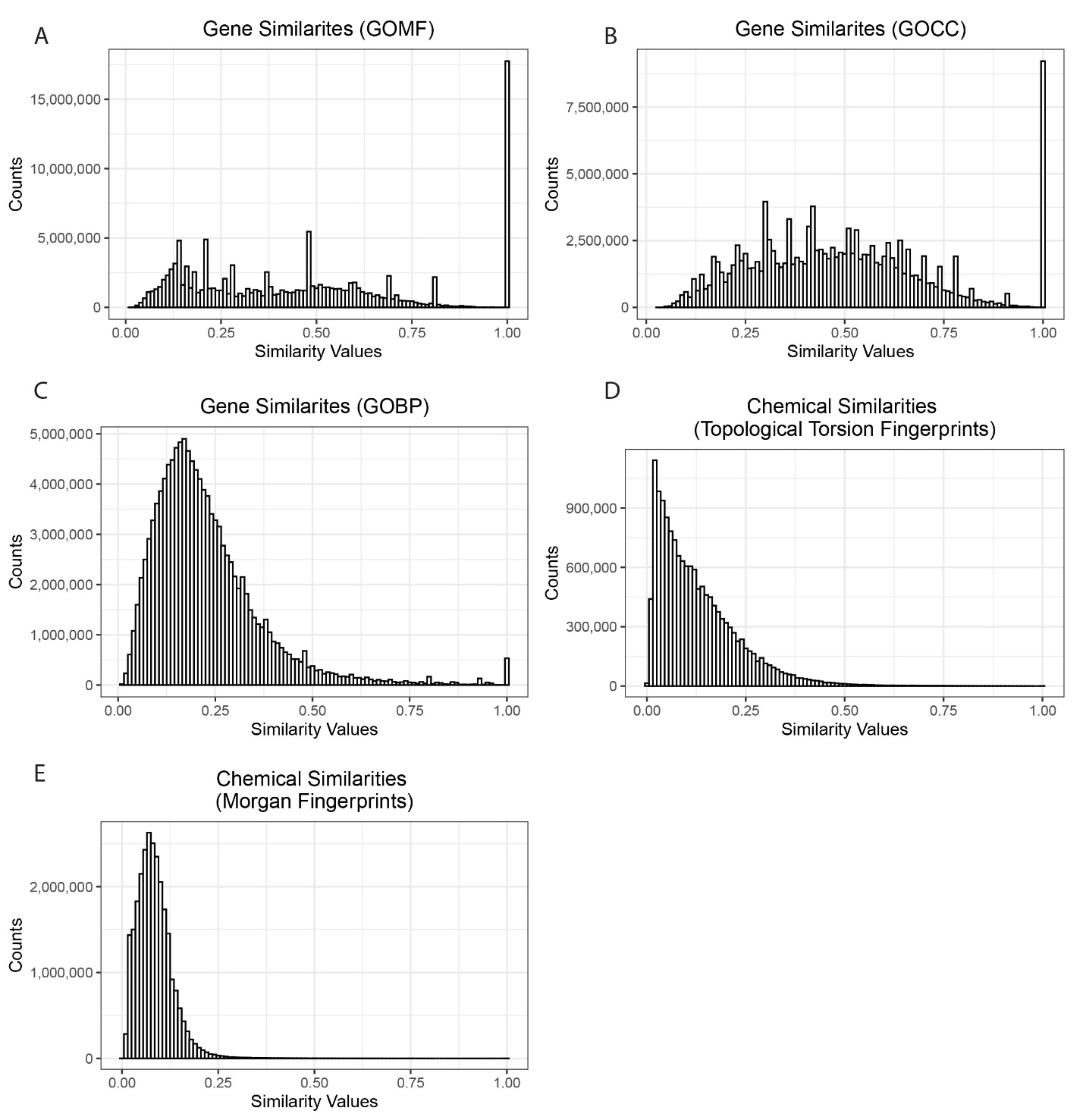

Supplementary Figure 1. Distributions of overall chemical and gene similarities. (A) Gene similarities calculated based on GOMF. (B) Gene similarities calculated based on GOCC. (C) Gene similarities calculated based on GOBP. (D) Chemical similarities calculated based on Topological Torsion Fingerprints. (E) Chemicals similarities calculated based on Morgan Fingerprints. Results based on 22,606 genes and 4,864 chemicals.

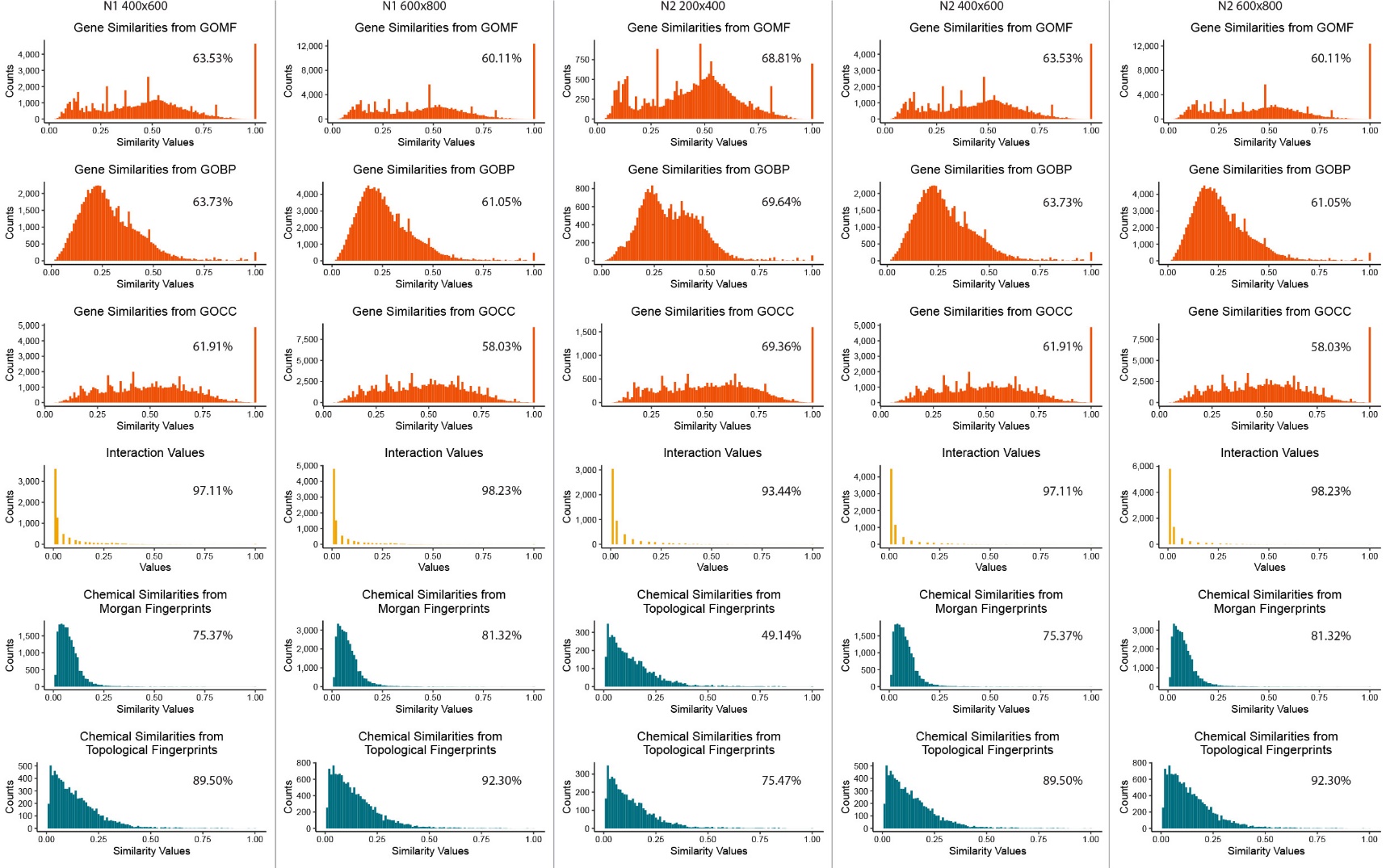

Supplementary Figure 2. Distributions of values in matrices used in the benchmark datasets. The sparsity of each matrix is labeled at the top right corner of each histogram.

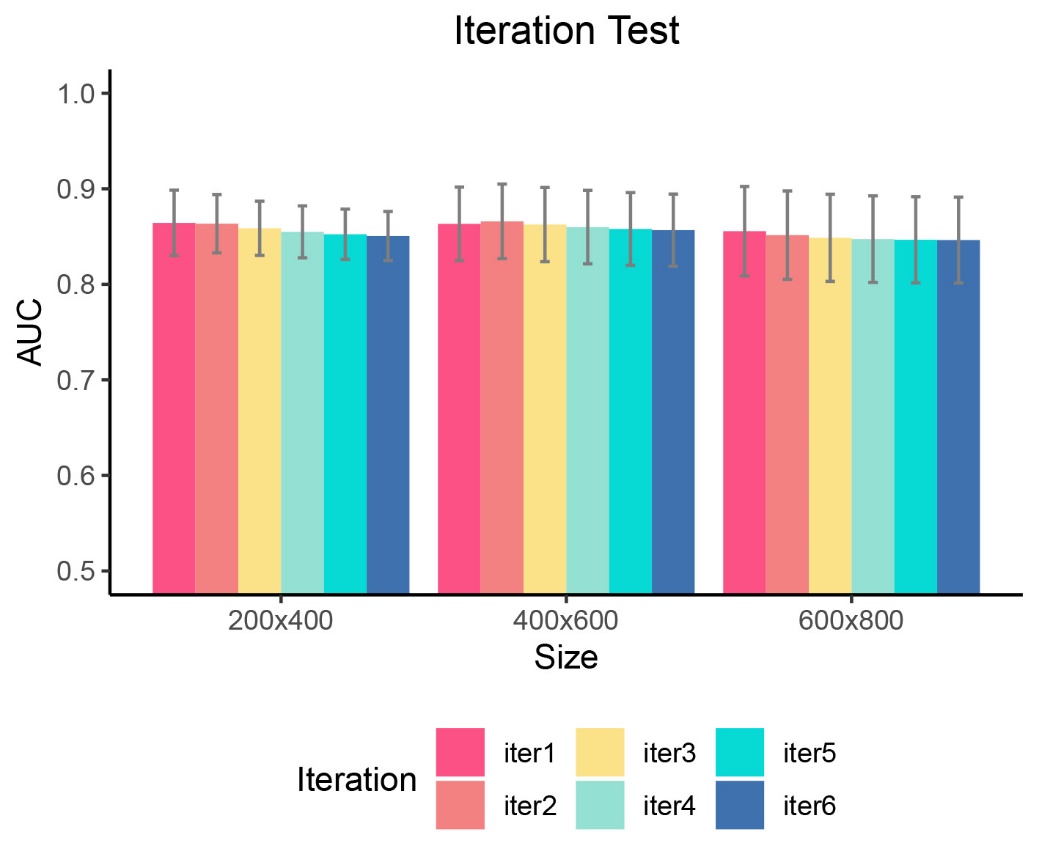

Supplementary Figure 3. The average AUCs of different iteration numbers. 0.9 was used as the hyperparameters for these calculations.

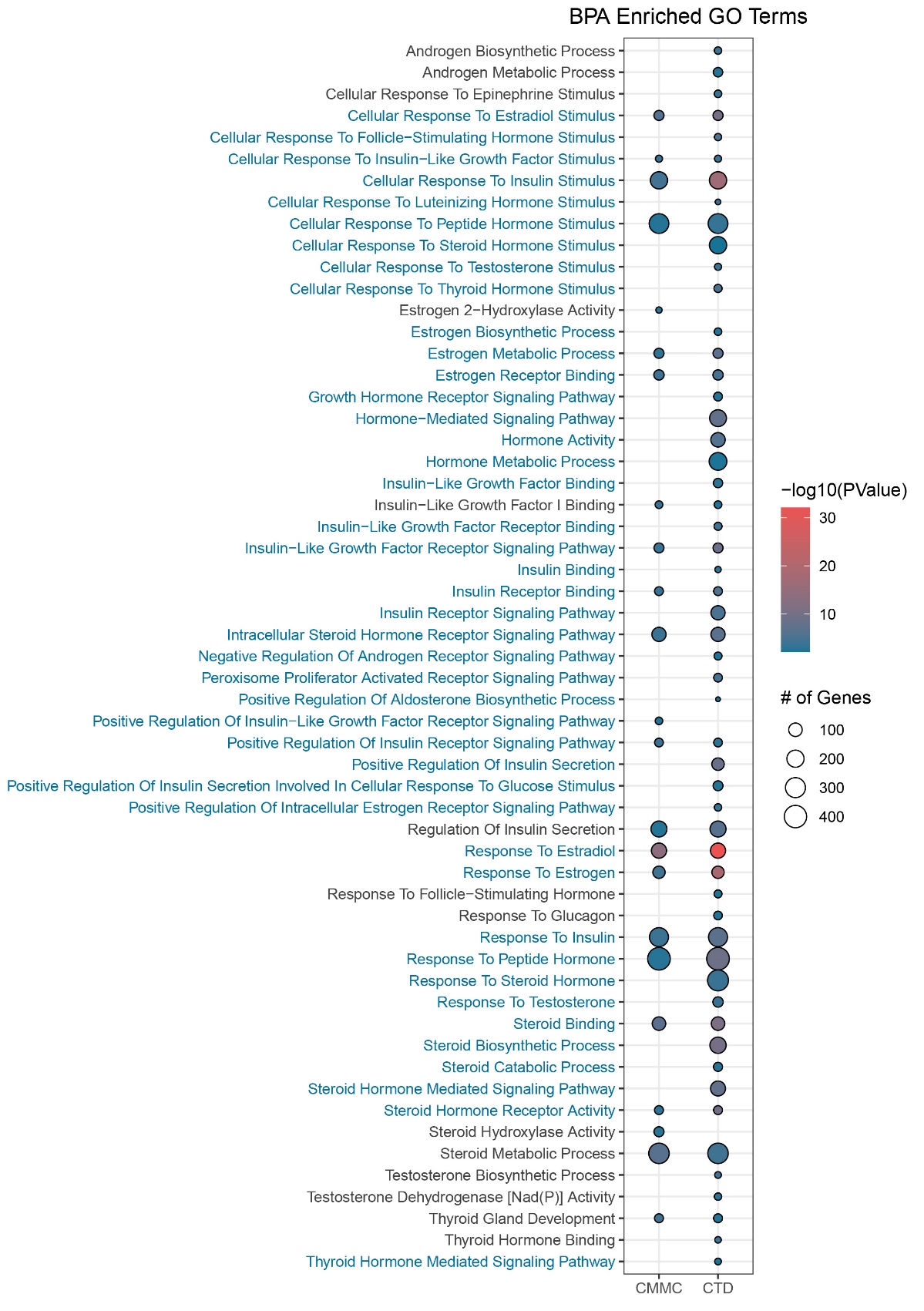

Supplementary Figure 4. Hormone-related significantly enriched GO terms from gene set enrichment test for BPA. Top 1000 interactive genes from our result and top 1000 interactive genes from CTD were used as input gene lists.

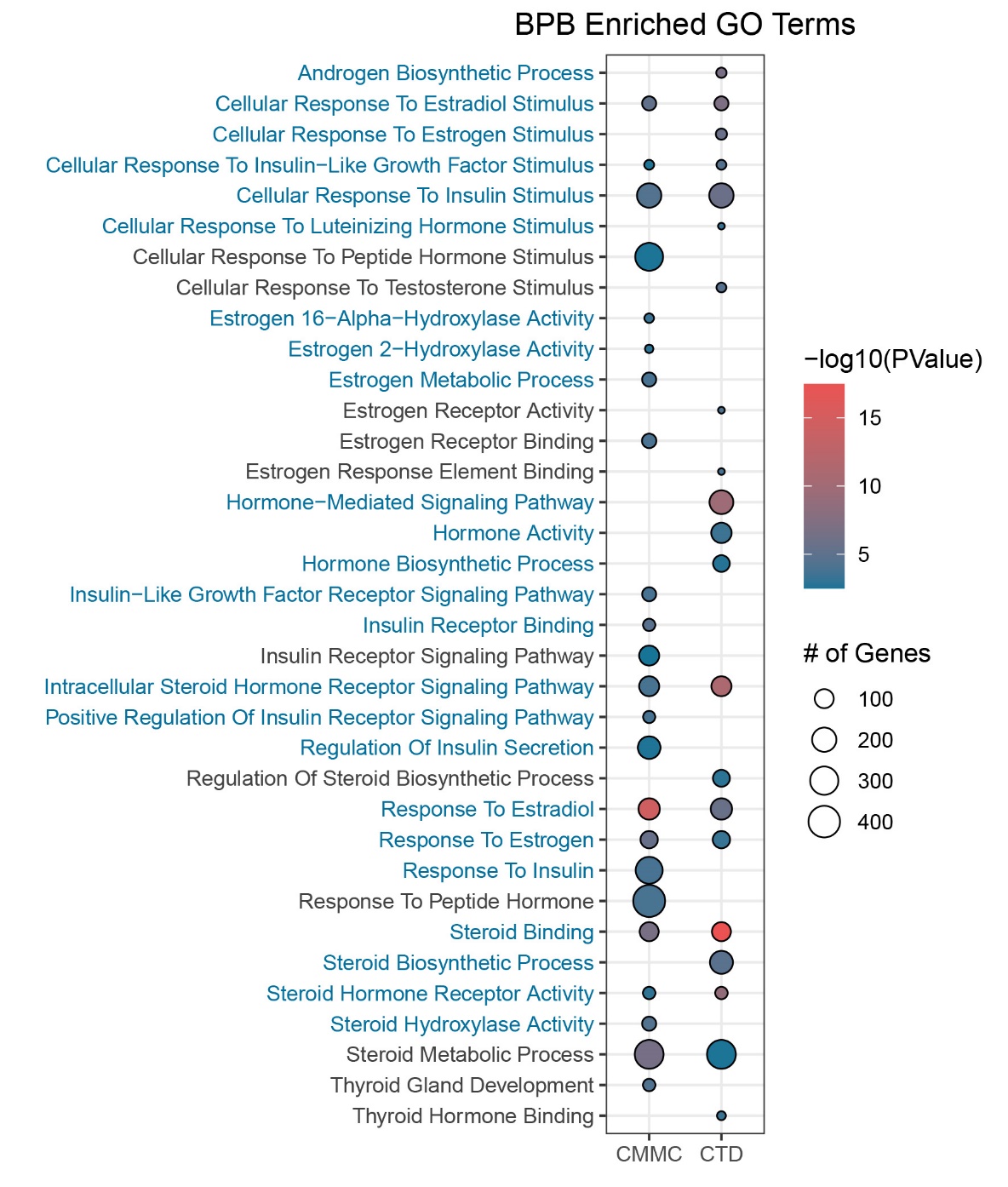

Supplementary Figure 5. Hormone-related significantly enriched GO terms from gene set enrichment test for BPB. Top 1000 interactive genes from our result and all 115 interactive genes from CTD were used as input gene lists.

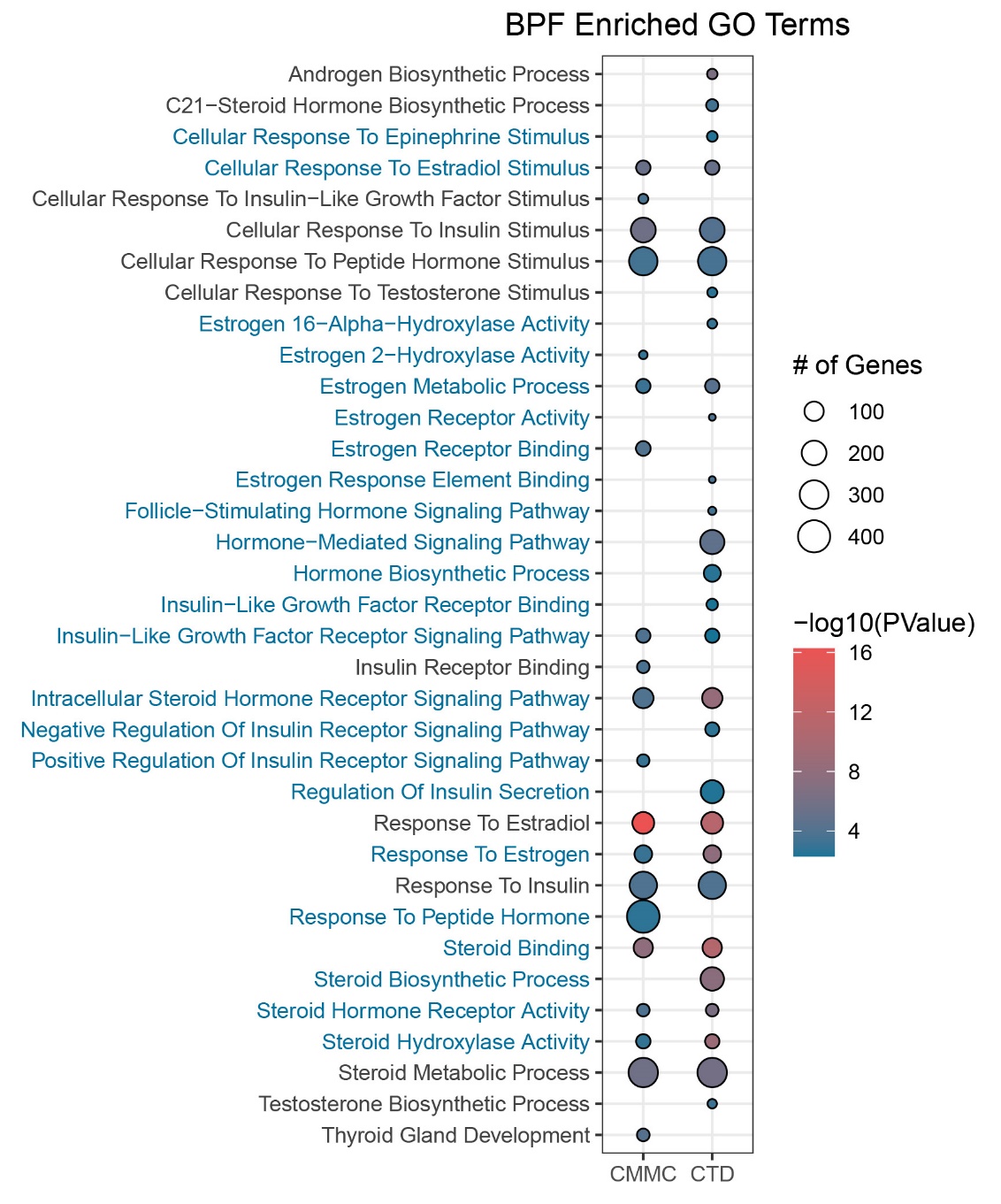

Supplementary Figure 6. Hormone-related significantly enriched GO terms from gene set enrichment test for BPF. Top 1000 interactive genes from our result and all 144 interactive genes from CTD were used as input gene lists.

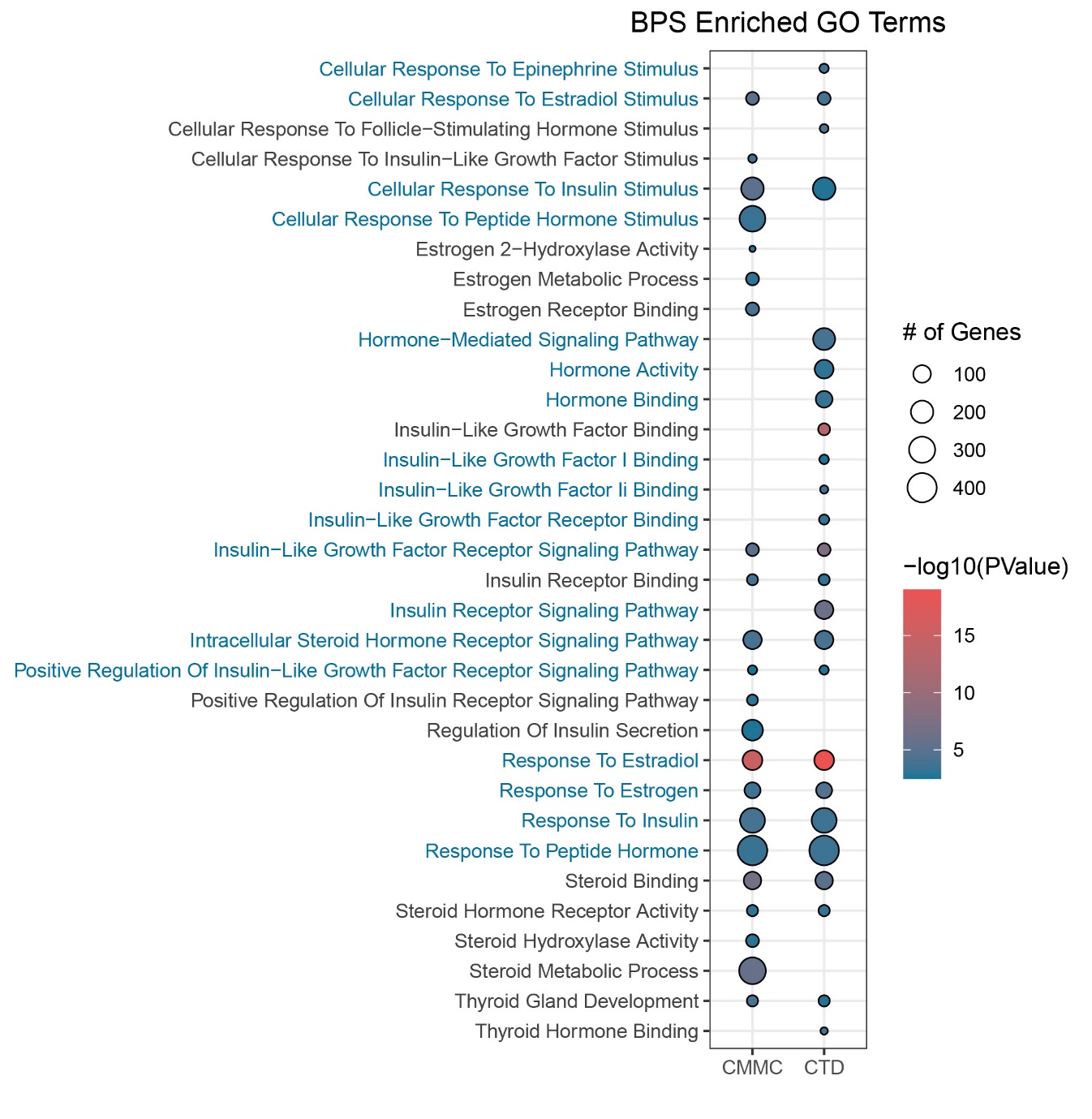

Supplementary Figure 7. Hormone-related significantly enriched GO terms from gene set enrichment test for BPS. Top 1000 interactive genes from our result and top 1000 interactive genes from CTD were used as input gene lists.

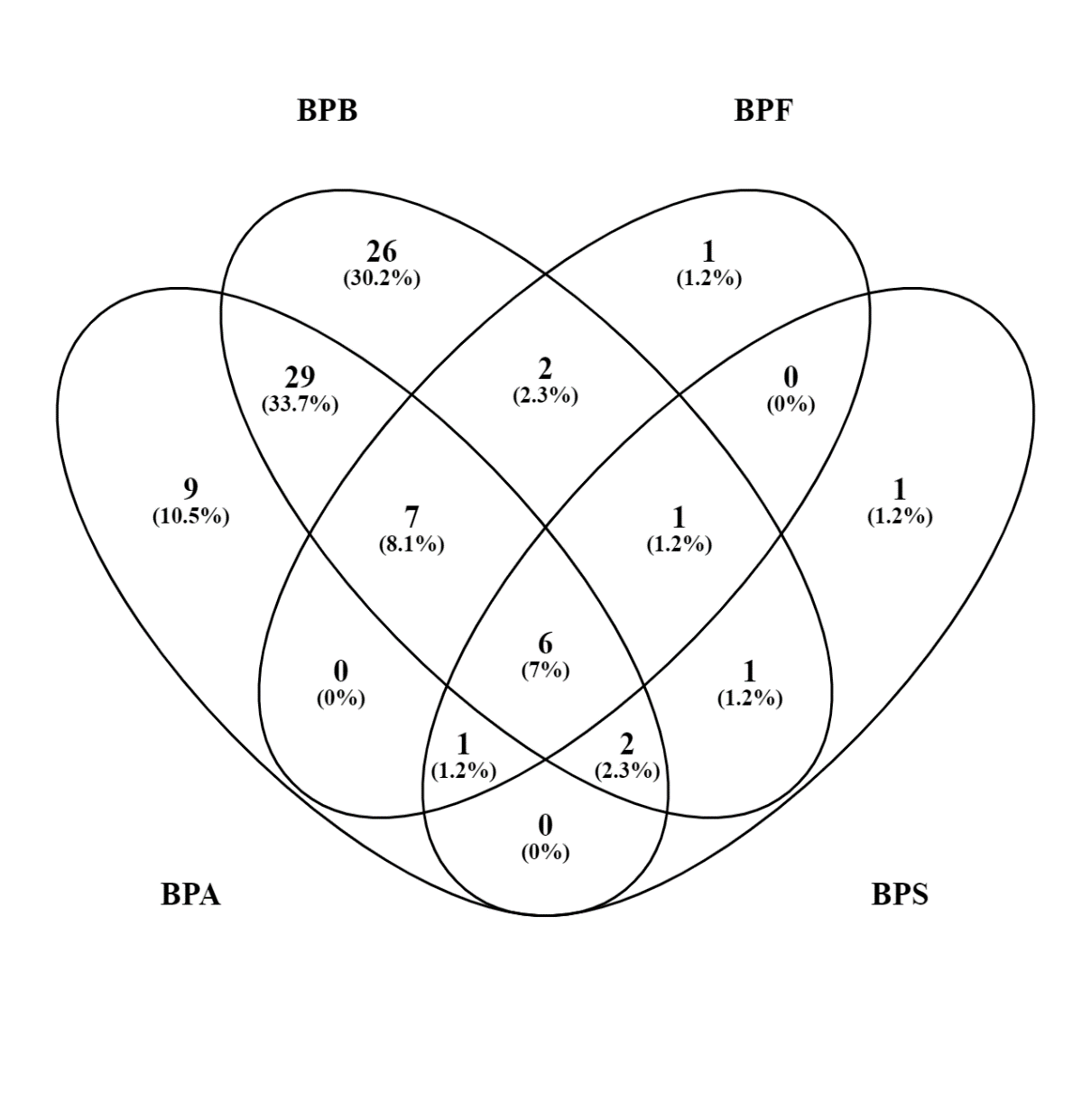

Supplementary Figure 8. CompTox active genes of these four chemicals used in our case study. *AHR* and *CYP3A4* are the two genes that predicted by CMMC as top target genes among the 9 BPA unique genes. *CCND1*, *CAT*, *NQO1*, *FOXO3*, *CXCL8*, *MMP9*, *NFIA*, and *ABCG2* are the eight genes that predicted by CMMC at top target genes among the 26 BPB unique genes. The unique gene of BPF and BPS was not predicted as top target genes by CMMC.

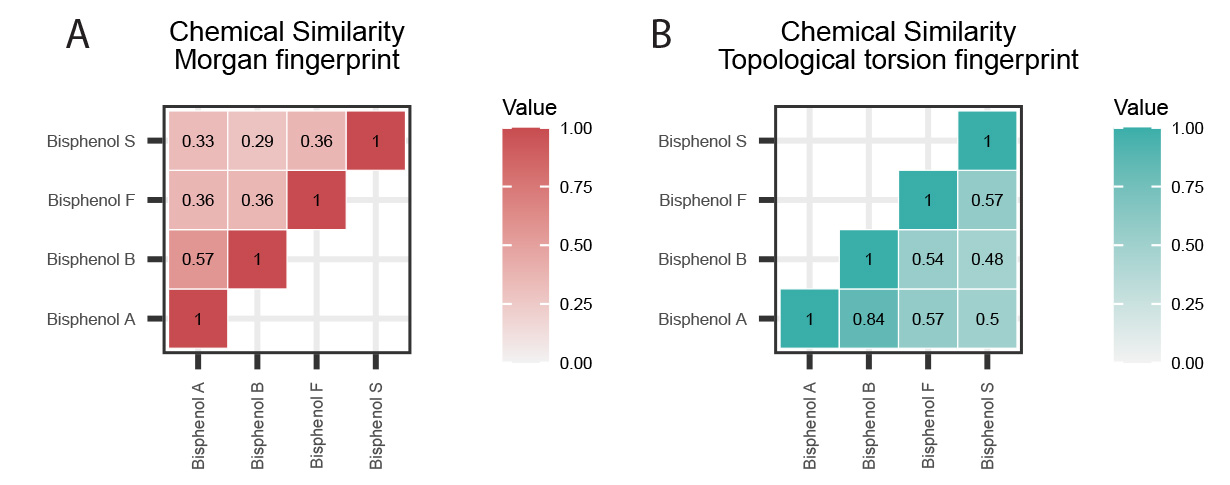

Supplementary Figure 9. Chemical similarities of BPA, BPB, BPF, and BPS. (A) Chemical similarities based on Morgan Fingerprints. (B) Chemical similarities based on Topological torsion Fingerprints.

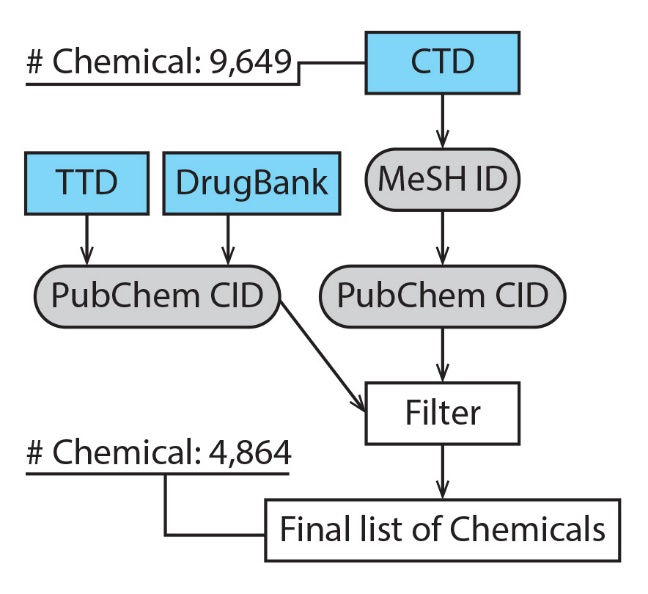

Supplementary Figure 10. Illustration of workflow for drug filtering. TTD and DrugBank were used to remove drugs included in the CTD dataset through PubChem IDs.

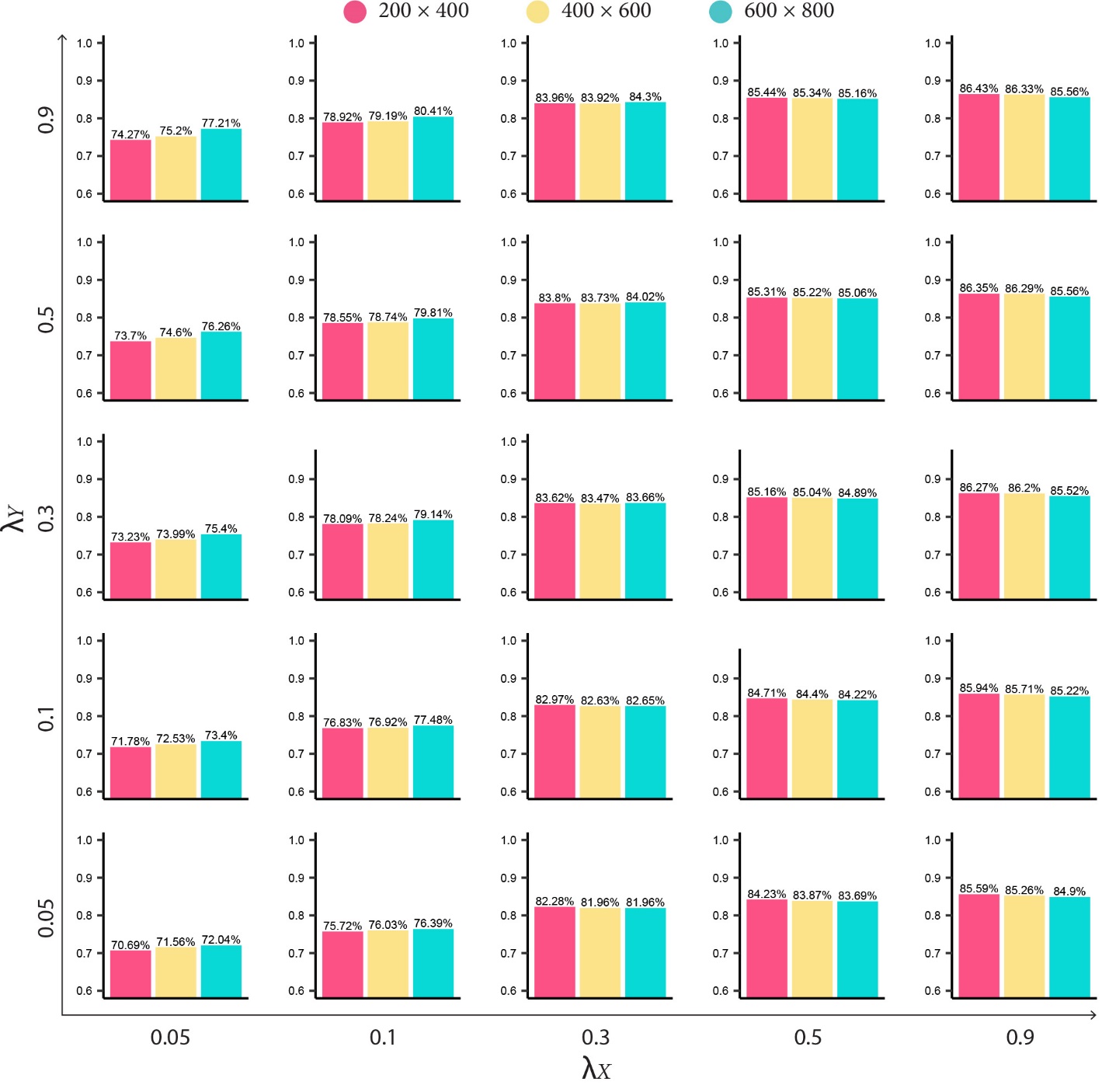

Supplementary Figure 11. Grid search results of hyperparameter tunning. Average AUCs across datasets are illustrated by the bar graphs.

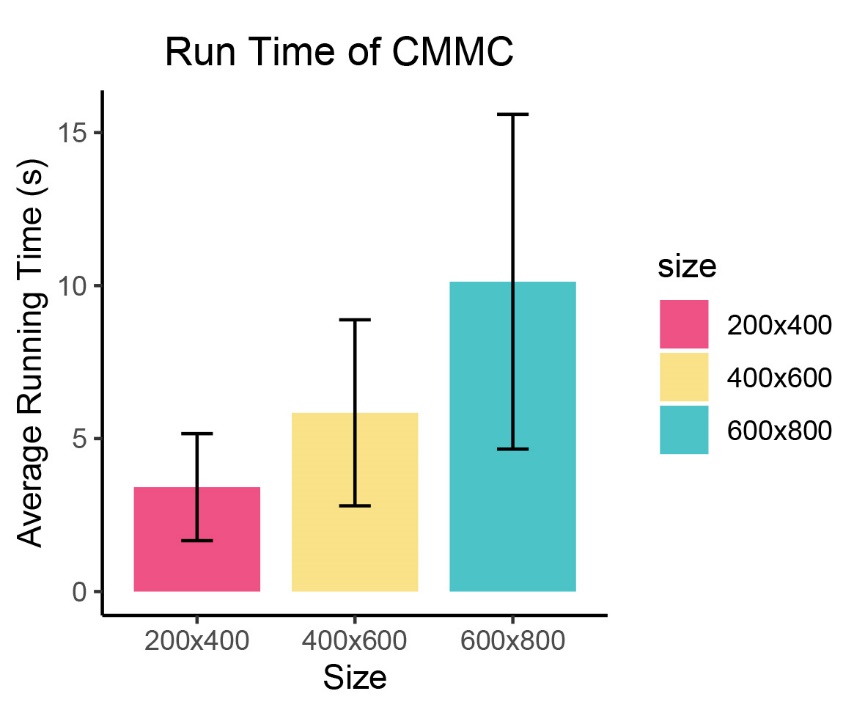

Supplementary Figure 12. Run time of CMMC in seconds with different sizes of input data. We executed CMMC on a Red Hat Enterprise Linux Server (release 7.9) with an Intel(R) Xeon(R) Gold 6150 CPU (2.70GHz). No parallel programming was used in the implementation.

Supplementary Table 1. Normalized and adjusted chemical-gene interactions.

Supplementary Table 2. Data summary of gene-gene similarities and chemical-chemical similarities.

| Chemical Similarity | | | | | | |
| --- | --- | --- | --- | --- | --- | --- |
| Types | Min. | 1st Qu. | Median | Mean | 3rd Qu. | Max. |
| Morgen Fingerprints | 0.002801 | 0.052632 | 0.078947 | 0.084704 | 0.108108 | 1 |
| Topological Torsion Fingerprints | 0.001129 | 0.048193 | 0.101695 | 0.127145 | 0.178218 | 1 |
| Gene Similarity | | | | | | |
| Types | Min. | 1st Qu. | Median | Mean | 3rd Qu. | Max. |
| GOBP | 0.007 | 0.134 | 0.198 | 0.2283 | 0.285 | 1 |
| GOMF | 0.015 | 0.207 | 0.423 | 0.4581 | 0.633 | 1 |
| GOCC | 0.027 | 0.314 | 0.471 | 0.4921 | 0.63 | 1 |

Supplementary Table 3. Performance comparisons on benchmark datasets with different binary conversion cutoffs. The best AUCs among different sizes of the input data were bolded. Our original plan was to use three additional cutoffs: 0.3, 0.4, and 0.5. However, because the interaction data is highly skewed towards the left, using 0.3, 0.4, or 0.5 as the binary conversion cutoff, resulted in too few “known neighbors” for WKNKN to estimate the likelihood of unknown interactions which caused an internal error. Thus, we only used the three cutoffs for performance comparisons.

Supplementary Table 4. Imputation results of CMMC for CompTox interactive genes. Separate tabs are used for BPA, BPB, BPF, and BPS.

Supplementary Table 5. Significantly enriched GO terms of BPA, BPB, BPF, and BPS. Input gene lists were the top 1000 ranked interactive genes from CMMC.

Supplementary Table 6. Total gene list and chemical list used in the case study.
